# Species turnover between age groups of horses and positive network of co-occurrences define the structure of horse strongylid communities: a meta-analysis

**DOI:** 10.1101/2021.05.17.441725

**Authors:** Michel Boisseau, Núria Mach, Marta Basiaga, Sławomir Kornaś, Tetiana Kuzmina, Claire Laugier, Guillaume Sallé

## Abstract

Grazing horses are infected by a wide range of strongylid species mostly located in the large intestine. Despite their impact on equine health and the emergence of drug resistant isolates, the phenology of these nematodes has been poorly characterized and the rules structuring their assembly as a community are not understood. Here, we compiled data on 46 equine strongylid species collected worldwide at the regional or horse scales (upon deworming or after necropsy) to analyse their richness, diversity and associated factors of variation. Worldwide, twelve species from the *Cylicocyclus* (n = 4), *Cylicostephanus* (n = 3), *Coronocyclus* (n = 2) and *Cyathostomum* (n = 2) genera were found in at least 75% of sites. Geoclimatic conditions had a limited effect on strongylid communities, but reduced species richness was found under the temperate European area. The recovery method did not affect species richness and differences on the temporal and sampling effort scales between studies applying either methods underpinned heterogeneous variances in community diversity.

At the horse level, rarefaction curves correlated poorly to parasite egg excretion, suggesting little contribution of community diversity to this trait. Using a diversity partitioning approach, we found that within-host diversity represented half of overall diversity underscoring the importance of host density and environmental contamination to the diversity of strongylid communities. While this is expected to erase diversity across communities, species turnover between age classes was the second most important contributor to overall diversity (23.9%). This was associated with a network of positive co-occurrences between the four most prevalent genera that we resolved at the anatomical niche level. Altogether this pattern of β-diversity maintenance across age classes combined with positive co-occurrences may be grounded by priority effects between major species. Our findings set the first assembly rules of equine strongylid communities.

## 1. Introduction

Grazing horses are naturally infected by a diverse parasitic fauna including Strongylidae (Taylor et al., 2012). They encompass subfamilies Cyathostominae and Strongylinae (Lichtenfels et al., 2008). The use of anthelmintics has resulted in prevalence levels for *Strongylus* sp. that has the highest fatality rate among Strongylinae (Herd, 1990), but has led to the emergence of resistant cyathostomin isolates worldwide (Matthews, 2014). affects horse growth and the massive emergence of encysted larvae can lead to fatal cases of cyathostomosis, the main cause of parasite mortality in young horses in Normandy, France (Sallé et al., 2020). Despite their importance in veterinary medicine, the phenology and the rules structuring the assembly of remain largely undefined due in part to the complexity of collecting data from large mammalian hosts under experimental conditions.

Inter-individual variation exists in host susceptibility to infection, with significant variation across age groups (Debeffe et al., 2016; Kornaś et al., 2015; Relf et al., 2013; Wood et al., 2013). In addition, climatic conditions affect the development of the free-living stages on pastures (Nielsen et al., 2007) and species reappearance following anthelmintic drug treatment varies across species (Kooyman et al., 2016) suggesting that combinations of their phenology or genetic diversity increase their fitness. Despite knowledge, variations in community structure in anatomical niches or environmental conditions remain largely uncharacterised.

Better defining the patterns of variation in helminth communities could however greatly improve their management in the field, i.e. by preventing the emergence of subdominant species following elimination of more dominant ones (Lello et al., 2004) or anticipating consequences of infections by other pathogens (Brosschot et al., 2021; Sweeny et al., 2020; Telfer et al., 2010). Data on helminth community assembly have been gathered across a wide range of host species including humans (Tchuem Tchuenté et al., 2003), rabbit (Lello et al., 2004), rodent (Behnke, 2008; Dallas et al., 2019), fishes (Poulin and Valtonen, 2002) with evidence of significant interactions between parasites. In horses, community structure has been determined from necropsy reports or after deworming(Gawor, 1995; Kornas et al., 2011; Kuzmina et al., 2005; Kuzmina et al., 2011; Kuzmina et al., 2016; Lind et al., 2003; Mfitilodze and Hutchinson, 1990; Ogbourne, 1976; Reinemeyer et al., 1984). However, these reports remain often descriptive and limited exploration of the relationship between community structure and host and environmental variables been conducted. For instance, variation in species richness and diversity along the intestinal tract of horses has been reported (Collobert-Laugier et al., 2002; Gawor, 1995; Ogbourne, 1976) and recent meta-analysis reported the impact of parasite recovery method and geographical region on cyathostomin abundance or prevalence (Bellaw and Nielsen, 2020). These studies however did not address how community structure, richness and diversity varied with environmental factors of interest. In addition, co-occurrence patterns between equine parasite species remain poorly characterized with contradictory findings reported (Sallé et al., 2018; Stancampiano et al., 2010).

Here, we performed an exhaustive literature search to evaluate how equine strongylid communities were shaped by environmental and host factors. In addition, we re-analyzed community data gathered from individual horses to partition diversity across environmental factors and to build co-occurrence networks at the anatomical niche level. Altogether, our results expand previous reports while opening new perspectives on the parasite composition and structure.

## 2. Materials and methods

The python and R scripts used to collect and analyse the data and associated datasets will be deposited at: https://github.com/MichelBoisseau37/script_meta_analyse upon manuscript acceptance.

### 2.1. Literature search and inclusion criteria

Studies were gathered from the Medline database (https://pubmed.ncbi.nlm.nih.gov/) using “strongyl* horse” as keywords, yielding a database of 1,032 articles. Article titles and abstracts were subsequently screened for particular terms including all possible combinations of the following: “intensity”, “Intensity”, “Abundance”, “abundance”, “Prevalence”, “prevalence”, “communit”, “Communit”, “Helminth” or “helminth”. To identify studies reporting strongylid burden, we also included the main genera (“*Cylicocyclus*”, “*Cyathostomum*”, “*Cylicostephanus*”) as keywords. This filtering step retained 300 papers. For these, titles and abstracts were read to determine whether findings of strongylid species prevalence and abundance would be available.

To broaden our database, we also run manual search on google (https://scholar.google.com/) using the keywords “prevalence abundance Cyathostominae horses” and “prevalence abundance Strongylidae horse”. This query identified six additional studies. Additional studies were identified after inspection of the references cited in three papers (Gawor, 1995; Ogbourne, 1976) that yielded prevalence data from 13 additional studies.

To be included in the meta-analysis, reported articles had to meet several criteria. First, studies were included in the dataset if they reported parasite prevalence (proportion of infected host), abundance (the number of parasite individuals per host, including uninfected individuals) or relative abundance. Second, published data were only included if the number of horses sampled was reported.

Finally, 39 papers met the aforementioned selection criteria (Supplementary Table 1). Among these, 16 papers presented both abundance and prevalence data, while others either only reported abundance (n = 5) or prevalence (n = 18) data. Papers from Africa (n = 2) and Australia (n = 3) were included to determine average relative abundance and prevalence across region-based studies but were not considered in any further analyses as too few replicates were available to test for environmental effects. For analytical purposes, papers were further divided into studies to account for the distinct experimental groups considered. This process yielded 69 community sets for 46 species encountered in at least one study.

### 2.2. Parasite, environmental and host variables

When available, metadata related to environmental factors were collected. Horse age was not considered for the continent-wide study as only scarce and inaccurate data were available. The same applied for horse breed or management type (wild or managed horses) that were reported in three studies only. Following edition, information regarding continent, country, sampling effort (number of horses sampled), the recovery method used (necropsy or deworming) were included.

Climate conditions were determined using the Köppen-Geiger climate classification (http://hanschen.org/koppen/#data) matching the sampling area. The Köppen-Geiger climate classification is encoded by a three- or two-letter system corresponding to the type of climate, rainfall pattern and temperature range respectively. For studies with no precise location indicated, the closest main city (state or country capital city) coordinates were used as a proxy (Supplementary Table 1). The different climates were clustered into broad climatic categories, namely continental, tropical, temperate and arid.

### 2.3. Statistical analysis

Data were analysed with R v. 3.6.3 and v. 4.0.2.

#### 2.3.1. Analysis of worldwide strongylid community diversity

##### 2.3.1.1. Considered regional community sets

Prevalence and relative abundance matrices were merged together into a single presence/absence matrix (noted 0 and 1 for absence and presence respectively). Analyses were run on the species presence/absence matrix for two reasons. First, this approach makes use of every available study (no matter of the reported measure) thereby providing greater statistical power than an approach considering abundance or prevalence data only. Second, most studies (*n* = 29 out of the 39 considered) did not report any dispersion parameter of parasite abundance and prevalence preventing any implementation of meta-analysis frameworks. In addition, because count data usually follow a negative binomial distribution, reported arithmetic mean values can be poor descriptors of the parasite population. This also speaks against sound estimation.

Because of the data structure, geoclimatic conditions (aggregated continent and climate zone) and the recovery method (deworming or necropsy) were confounded. In addition, some conditions were under-represented (less than five community sets) and not considered further. This included the community sets from Asia (n = 1), Australia (n = 3), or sets from temperate Europe collected after deworming (n = 3).

As the deworming strategy was only applied in Europe, a continental European subset was considered (*n* = 33 communities) to evaluate how the recovery method could affect the strongylid diversity. Similarly, the geoclimatic zone effect was tested on communities recovered under the same necropsy framework leaving 30 community sets that fell in temperate America (n = 8), tropical America (n = 7), temperate Europe (n = 11) or continental Europe (n = 5).

##### 2.3.1.2. Diversity analyses

Species richness was taken as an α-diversity estimator, and the β-diversity index was estimated with the Jaccard index using the vegan R package. (version2.5-7). Inter-sample distances were visualized using the Non-Metric Scaling (NMDS) ordination method through the “metaMDS” function in the vegan R package. This method generated fictitious axes maximizing the variance and allowing an optimal visualization of distances. NMDS is the most appropriate representation because it minimizes dissimilarities between similar communities and vice versa for more remote communities. NMDS was run on the whole set of communities to establish how environmental factors (aggregated factor of the geoclimatic zone and the recovery method), the sampling effort (number of horses) and the year of publication were related to community structures.

A permutational multivariate analysis of variance (PERMANOVA) using distance matrix (adonis function of the vegan R package) was then applied to identify the factors that would influence most the community structure and to obtain the percentage of variance explained by the factors of interested, i.e. sampling effort (number of horses), the geoclimatic zone and the recovery method using appropriate subsets to control for environmental conditions.

The respective effects of the recovery method (tested on the European subset) or the geoclimatic area (tested on the necropsy-based subset) on species richness were estimated using a linear model, fitting sample size, year of publication and the factor of interest as fixed effects.

##### 2.3.1.3. Effect of environmental factors on parasite prevalence across region-based community sets

Last, we aimed to identify strongylid species whose presence or absence in a given set would be associated with variation in the recovery method or the geoclimatic area. This was applied to the aforementioned community subsets to account for the data structure. In each case, the presence or absence of worm strongylid species was modelled using a logistic regression model, fitting an interaction term between worm species and either the recovery method (European continental subset) or the geoclimatic area (necropsy-based data). Because quasi-complete separation of species with environmental factors of interest occurred (identified with the “detect_separation” function of the *detectseparation* v0.1 package), we restricted this analysis to ten species. In addition, model convergence could not be achieved in case the community set was fitted as a random effect. To deal with this while accounting for inter-sets variation, we fitted the year of publication, sampling effort and the species richness of the community as covariates.

#### 2.3.2. Analysis of diversity at the horse level

These analyses were run on horse-based strongylid community data from three published studies, either collected after the necropsy of 46 horses in France (Collobert-Laugier et al., 2002) or the deworming of 48 horses in Poland (Sallé et al., 2018) and or 197 horses Ukraine (Kuzmina et al., 2016) respectively.

##### 2.3.2.1. Relationship between diversity and faecal egg count at the horse level

To evaluate how strongylid community diversity could relate to measured Faecal Egg Count (FEC) in their host, we estimated Spearman’s correlations between FEC and species richness, or FEC and the Gini-Simpson index which corresponds to the slope of the species accumulation curve at its basis and relates to species dominance (Chase and Knight, 2013). This was not done on the necropsy data as FEC measures were not available.

##### 2.3.2.2. Impact of horse age on strongylid community diversity

To study the interactions between strongylid species within their hosts, we re-analyzed individual necropsy report data collected in Normandy (Collobert-Laugier et al., 2002). Abundances of 20 strongylid species and other non-strongylid species (*Anoplocephala* sp., *Oxyuris sp*.) were quantified within the caecum, ventral- and dorsal-colon of 46 horses. The data from 36 horses that had their age registered and were infected by at least one parasite species were kept. To avoid spurious signals associated with rare observations, we focused on the 15 species achieving at least 10% prevalence. To capture the strongylid community assembly pattern across anatomical niches and age group, we applied the hierarchical Bayesian joint species distribution model (Ovaskainen et al., 2017; Tikhonov et al., 2017) implemented in the HMSC R package (v. 3.0-10) (Tikhonov et al., 2020). This model offers an integrated framework to explicitly model species-species interactions while simultaneously accounting for environmental covariates, species traits or their phylogenetic relationships (Ovaskainen et al., 2017; Tikhonov et al., 2017). We used the same approach as described elsewhere (Abrego et al., 2020), fitting two models that either accounted for the worm burden measured at the horse level or for additional fixed effects including the anatomical niche and horse age. In both cases, horse and horse *x* anatomical niche were fitted as random effects to account for inter-horse variation in species pairs co-occurrences and for sample level variation not accounted for otherwise (Abrego et al., 2020). A hurdle-type model was considered to account for the overdispersed nature of the count data, modeling log-transformed positive species counts as normal and the binary presence or absence status using a probit-link function (Abrego et al., 2020). Model predictive power was assessed by Area Under the Curve (AUC) and R^2^ parameters averaged across species for presence-absence and abundance data. Parameters were estimated from 4,000 posterior samples collected every 100 iterations from four Markov Chain Monte Carlo (MCMC) run for 150,000 iterations with a third discarded as burn-in.

#### 2.3.3. Analysis at the anatomical niche level: species co-occurrences from necropsy data

To study the interactions between strongylid species within their hosts, we re-analyzed individual necropsy report data collected in Normandy (Collobert-Laugier et al., 2002). Abundances of 20 strongylid species and other non-strongylid species (*Anoplocephala* sp., *Oxyuris sp*.) were quantified within the caecum, ventral- and dorsal-colon of 46 horses. The data from 36 horses that had their age registered and were infected by at least one parasite species were kept. To avoid spurious signals associated with rare observations, we focused on the 15 species achieving at least 10% prevalence. To capture the strongylid community assembly pattern across anatomical niches and age group, we applied the hierarchical Bayesian joint species distribution model (Ovaskainen et al., 2017; Tikhonov et al., 2017) implemented in the HMSC R package (v. 3.0-10) (Tikhonov et al., 2020). This model offers an integrated framework to explicitly model species-species interactions while simultaneously accounting for environmental covariates, species traits or their phylogenetic relationships (Ovaskainen et al., 2017; Tikhonov et al., 2017). We used the same approach as described elsewhere (Abrego et al., 2020), fitting two models that either accounted for the worm burden measured at the horse level or for additional fixed effects including the anatomical niche and horse age. In both cases, horse and horse *x* anatomical niche were fitted as random effects to account for inter-horse variation in species pairs co-occurrences and for sample level variation not accounted for otherwise (Abrego et al., 2020). A hurdle-type model was considered to account for the overdispersed nature of the count data, modeling log-transformed positive species counts as normal and the binary presence or absence status using a probit-link function (Abrego et al., 2020). Model predictive power was assessed by Area Under the Curve (AUC) and R^2^ parameters averaged across species for presence-absence and abundance data. Parameters were estimated from 4,000 posterior samples collected every 100 iterations from four Markov Chain Monte Carlo (MCMC) run for 150,000 iterations with a third discarded as burn-in.

## 3. Results

### 3.1. Descriptive features of strongylid communities

The breakdown of publication counts for factors of interest is provided in Supplementary Table 2. As can be seen in Figure 1A, studies using deworming were mostly carried out in Eastern Europe. Our data showed an average number of horses per study of 28.65, ranging between 2 and 150 individuals. Strongylid communities were dominated by a small number of species, *i*.*e. Cylicocyclus nassatus, Cylicostephanus longibursatus, Cyathostomum catinatum* (Figure 1B, supplementary table 3) with an average prevalence of 83.97% ± 3.38. Along these three dominant species, a set of eight subdominant species with an average prevalence of 54.96% ± 12.68. On the other hand, 31 rare species had an average prevalence of 8.93% ± 6.70. These mean prevalence levels reported within horse populations were slightly corroborated when considering the presence-absence pattern at the regional scale (Figure 1C). Specifically, the dominant and subdominant species defined by an average prevalence of 25% or more at the horse population level were present in 75% of studies. Contrasting this broad pattern, *Cylicostephanus goldi* was relatively frequent among horses but its presence varied widely across studies (Figure 1C).

**Figure 1.**
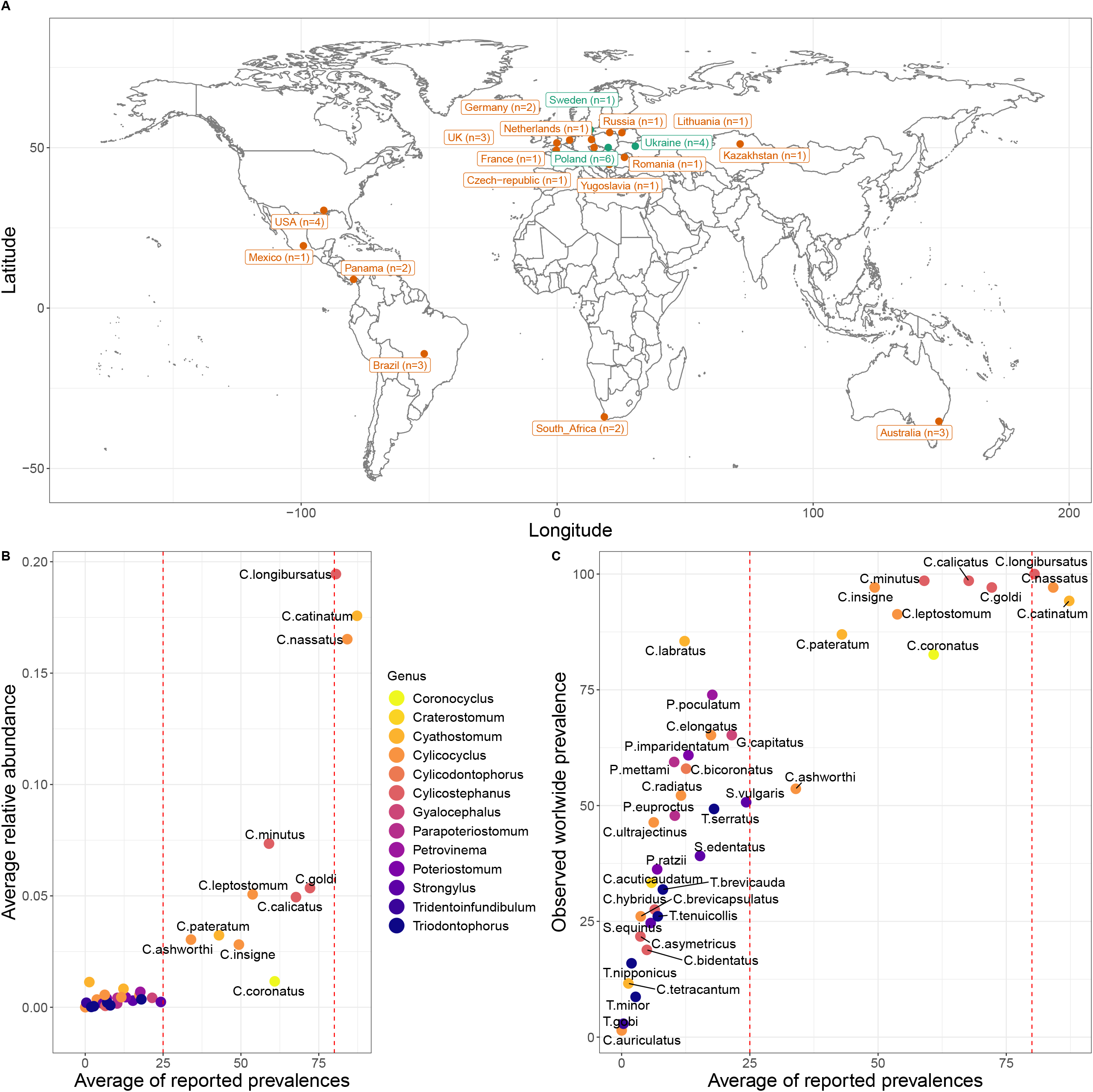
Descriptive features of worldwide equine strongylid communities. A: World map of study distribution coloured by worm collection method, either following deworming (green) or necropsy (orange). B: Strongylid species (colored by their respective genera) relative abundances and prevalences averaged across published mean estimates. Dotted lines distinguish between rare, subdominant and dominant species in the left, middle and right quadrants respectively. C: Comparison of observed species prevalence across studies (from the presence-absence matrix) against the average published prevalence estimates. Colour and dotted lines match that used in B.

At the horse level, strongylid community diversity varied widely as illustrated by species accumulation curves derived from the three available studies. The sampling of 95% of available diversity, required 22 horses (46% of total horse sample) for necropsy-based data from France (Collobert-Laugier et al., 2002). On the contrary, strongylid communities gathered upon deworming reached the same threshold for 25% of available horses (12 and 50 horses in the studies in Poland or Ukraine communities). More diverse strongylid communities may reflect reduced drug usage and might be associated with higher parasite egg excretion. However, limited agreement was found between Faecal Egg Count and community diversity as measured by the rarefaction curves. Significant correlation was found between FEC and species richness (Supplementary Figure 1) in strongylid communities gathered in Poland (Spearman’s *ρ* = 0.31, *P* = 9×10^−3^) but not in Ukraine (Spearman’s *ρ* = 0.03, *P* = 0.69). Correlation between FEC and the rarefaction curve at its basis (Gini index) was not significant in any case (Spearman’s *ρ* = -0.14, *P* = 0.06 and *ρ* = 0.24, *P* = 0.1 in Polish and Ukrainian data respectively), suggesting that species dominance does not affect FEC.

### 3.2. Diversity analysis of worldwide strongylid communities

As a first step, an exploratory multivariate analysis was applied to our meta-community matrix (Figure 2). This analysis highlighted the outstanding contribution of the strongylid recovery method on community diversity (Figure 2). This clustering also closely matched a gradient defined by differences in the temporal and the sampling effort scales. Publication times ranged between 2005 and 2018 or between 1935 and 2016 for deworming- and necropsy-based sets respectively. In addition, significantly more horses were sampled in necropsy-based studies (36.84 and 18.9 horses sampled respectively, *P* = 2.2 × 10^−16^).

**Figure 2.**
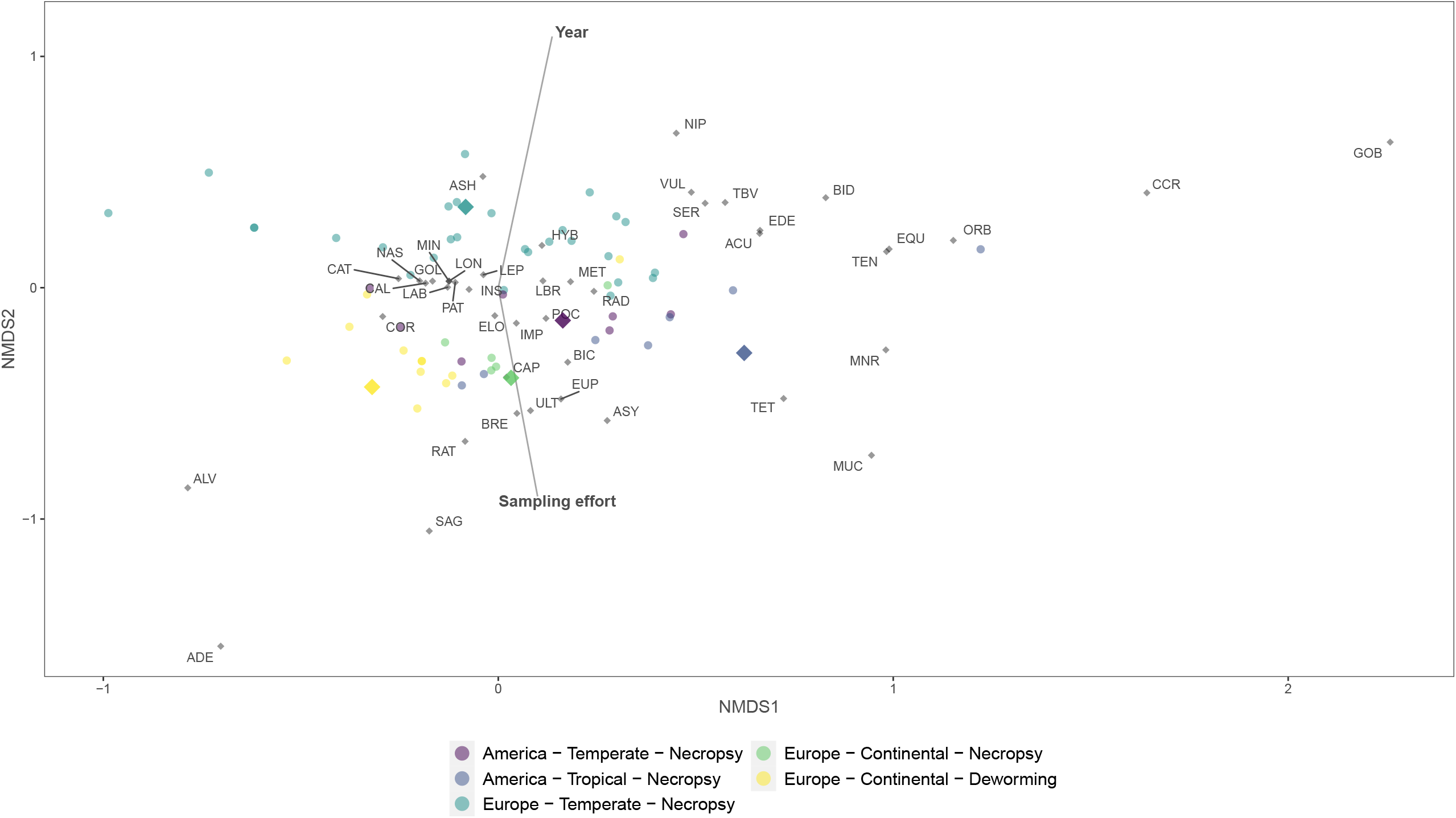
Ordination applied to strongylid meta-community with fitted environmental variables. Figure shows the first two axes from a NonMetric Dimensional Scaling (NMDS) ordination applied to published metacommunity data. The colours correspond to the different types of environment considered, coded as Continent – Climate – Recovery method. For visualization, species names were abbreviated with correspondence given in Supplementary Table 3.

The clear distinction between considered recovery methods was supported by the PERMANOVA (*P* = 3 × 10^−4^) run on the strongylid communities gathered under the same geo-climatic conditions, i.e. European continental climate. This was however associated with higher variance in deworming-based methods (F_1,31_ = 6.51, *P* = 0.02, supplementary Figure 2).

Logistic regression models found significant variation for *Triodontophorus serratus* and *Strongylus vulgaris* that were less often recovered in the five considered necropsy-based sets relative to the 28 deworming studies performed under the same geoclimatic conditions (interaction terms equal to -4.1 ± 1.96, *P*=0.03 and -4.62 ± 2.00, *P* = 0.02). Of note, quasi-complete separation with the recovery method was found for 13 species, either showing complete absence (*Cylicostephanus. bidentatus*) or complete presence (*Cylicocyclus ashworthi, Gyalocephalus capitatus, Cylicodontophorus bicoronatus, Coronocyclus coronatus, Cylicocyclus. elongatus, Coronocyclus labratus, Poteriostomum imparidentatum, Petrovinema poculatum, Parapoteriostomum mettami, Poteriostomum ratzi, Cylicocyclus Radiatus, Triodontophorus tenuicollis*) in studies practicing necropsy (Supplementary Figure 3). The species richness was however not significantly different between the recovery methods (*P* = 0.6).

On the contrary, the ordination analysis found limited clustering associated with geoclimatic factors (Figure 2). This was corroborated by PERMANOVA on the set of communities collected after necropsy (*P* = 1.9 × 10^−3^). In that case, species richness was lower in the temperate European area (−6.57, *P* = 5×10^−3^). Apart from the 15 species with quasi-complete separation in one of the considered geoclimatic zones (Supplementary Figure 4), we did not find significant evidence of variation in their prevalence pattern across geoclimatic conditions.

To sum up, the recovery method was not associated with variation in species richness but it was associated with significant turnover across communities, likely mirroring scale differences between the considered sets. On the contrary, species turnover across geoclimatic areas was limited and reduced richness was found under temperate Europe.

### 3.3. Strongylid community dynamic across age groups

Partitioning of overall *γ*-diversity across Ukraine operations revealed an outstanding contribution of within-horse α-diversity (46.5% of overall diversity) relative to species turnover coefficients (Figure 3). The observed value was however lower than under null model expectations (22.6% difference, *P*=10^−3^). Species turnover between horse age-group was the second most important contributor to *γ*-diversity (23.9% of *γ*-diversity), significantly higher than that expected from a null model (*P*=10^−3^, Figure 3) and higher than that explained by between-farm variation (20% of *γ*-diversity, Figure 3). Of note, observed between-horse turnover did not depart from the simulated null distribution (0.06% difference, *P*=0.4, Figure 3).

**Figure 3.**
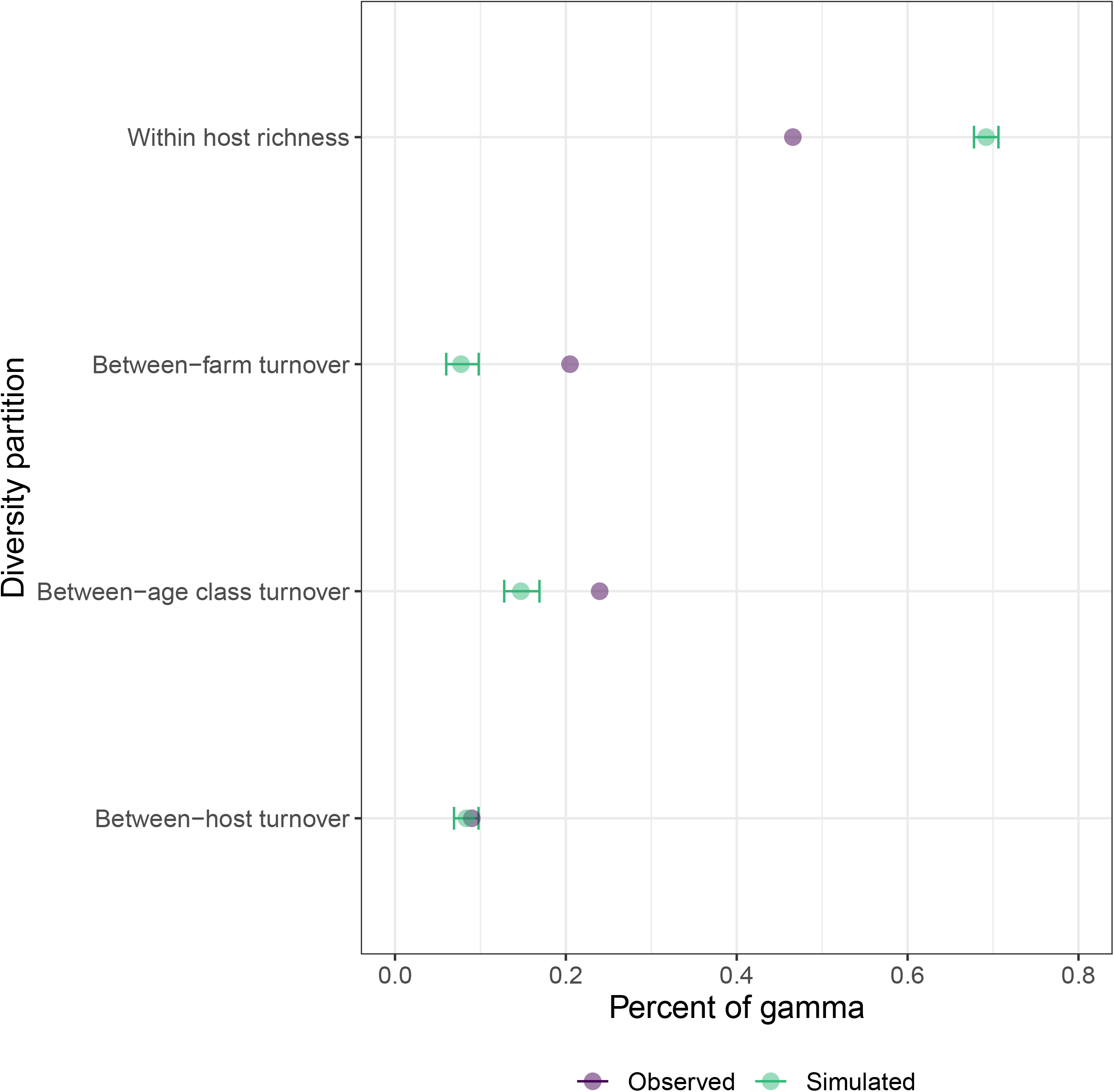
Hierarchical diversity partitioning of overall diversity. Within host alpha diversity and β-diversity (turnover) for the scales of interest (host, age class and farm), observed (purple) and simulated (green) estimates are given in percentage of overall diversity explained. Error bars materialize a 95% confidence interval of the simulated data.

Because of the importance of age group in driving strongylid community structure, we also searched for indicator species that would preferentially be associated with a given age group. This analysis found that *C. labiatus* and *C. ashworthi* were preferentially encountered in two-years old horses, irrespective of the considered coefficient (indicator values of 0.659, *P*=7 × 10^−3^, and 0.585, *P* = 0.04 or association coefficient equal to 0.42, *P* = 0.01 and 0.41, *P* = 0.01 for *C. labiatus* and *C. ashworthi*, respectively).

Altogether these results suggest that horse-level community diversity and between age-class turnover are the major contributors driving measured diversity under the European continental conditions encountered across Ukraine operations.

### 3.4. Co-occurrence of species within different anatomical niches

Anatomical niche was the last scale examined to determine factors driving strongylid community structure. The predictive power of the modeling approach was high with a mean AUC of 0.88 and R^2^ of 0.79 for models based on presence-absence or positive abundance data.

Raw or residual co-occurrences only supported positive interactions between the considered strongylid species (Figure 4). The correction for environmental covariates erased around half of observed co-occurrences in both abundance (11 out of 26) and presence-absence (21 out of 42) models. This was mostly due to correction for the anatomical niche that accounted for a third of total variance while another third was equally shared across horse age and infection intensity (Table 1). Of note, the presence of *C. goldi* was largely dependent on inter-horse variation but not its abundance (supplementary Figures 5 and 6) suggesting limited expansion abilities once established in a host. Residual models shared 12 positive co-occurrences involving *C. nassatus, C. labratus, C. coronatus, Cylicostephanus minutus* and *C. catinatum* (Figure 4). However, more co-occurrences were found with the presence-absence data. This suggests that strongylid species tend to have similar co-infection patterns but their simultaneous presence does not necessarily affect their respective relative abundances.

**Table 1.** Proportion of variance explained by considered covariates in the residual co-occurrence model. For each covariate of interest (fixed or random effect), the proportion of variance explained is given for the two model types considered using positive abundance data or presence-absence data.

| Effect type | Covariate | Abundance data (%) | Occurrence data (%) |
| --- | --- | --- | --- |
| Fixed effect | Anatomical niche | 31.2 | 32.7 |
|  | Horse age | 14.6 | 12.9 |
|  | Worm burden | 14.1 | 10.8 |
| Random effect | Horse | 12.8 | 33.4 |
|  | Horse x niche | 27.3 | 19.1 |

**Figure 4.**
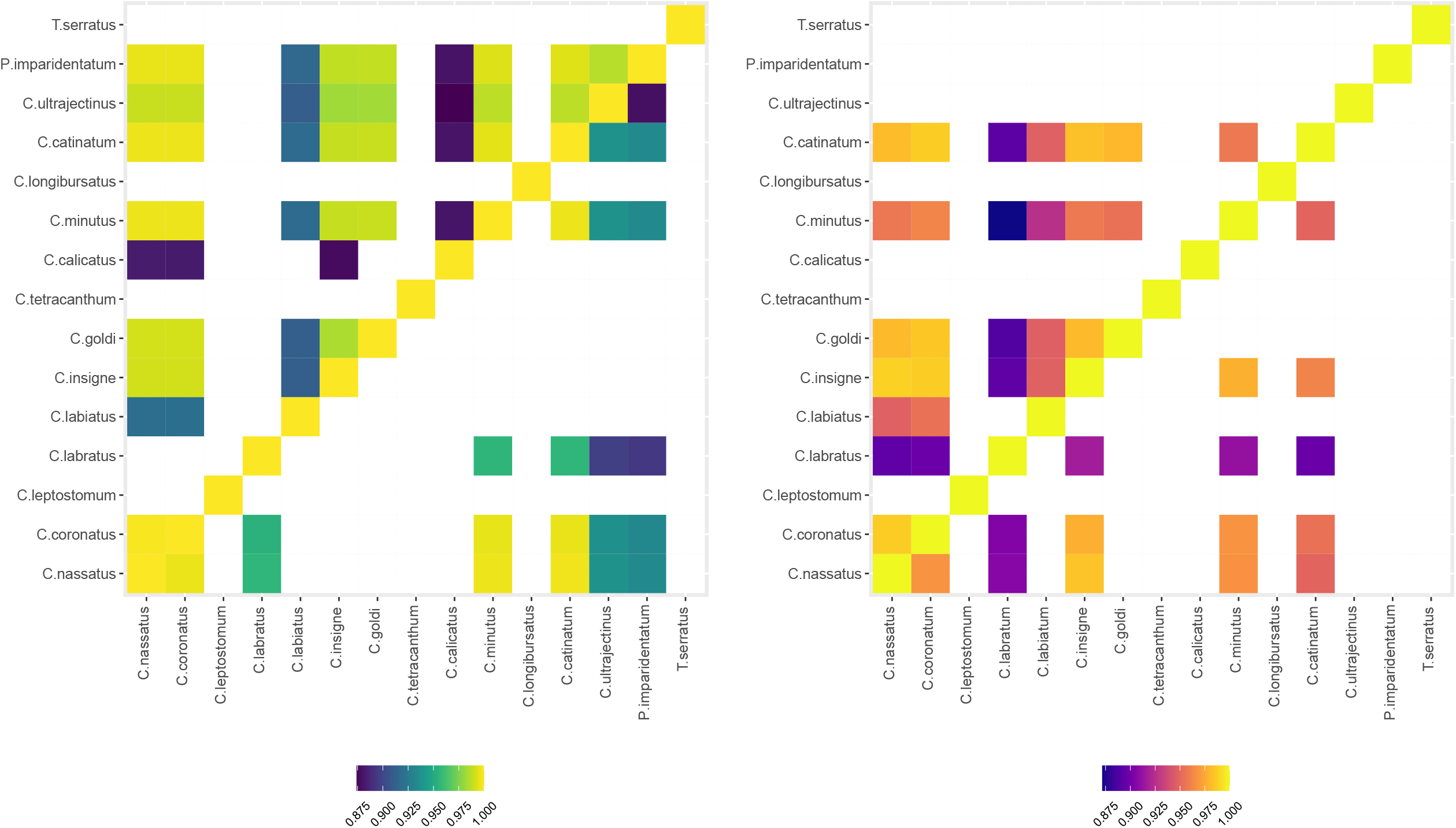
Raw and residual strongylid species co-occurrence matrices estimated from presence-absence data or positive abundance data. Co-occurrence matrices derived from presence-absence (left) or positive abundance (right) data are represented for 15 strongylid species collected identified in 36 necropsied horses from Normandy, France. In each case, off-diagonal elements describe the raw (above) and residual (below) co-occurrences.

## 4. Discussion

In this meta-analysis, we have explored the principles of strongylid community structuring across various scales, ranging from regional down to anatomical resolutions. Our findings define a community with little rearrangements across contrasted geoclimatic conditions, whose diversity is mostly grounded on within-host richness and turnover across-age groups. At the anatomical scale, strongylid communities are structured around positive co-occurrences fluctuating across anatomical niches. As such, our results complement and expand the past work by Bellaw and Nielsen (Bellaw and Nielsen, 2020) with additional horse-based data and different approaches and interpretations of the findings. While we tried to account for common sources of variation at the regional scale, this was often limited by the paucity of available metadata or by the data structure of past published work. For instance, age group was poorly annotated across studies and horse sex was seldom reported despite significant species turnover described between both sexes (Sallé et al., 2018). In addition, the lack of appropriate summary statistics in past published work constrained our analysis to a presence-absence framework that certainly overlooks finer grain patterns found with abundance data (Brian and Aldridge, 2020). This is mostly the result of the existing asymmetry between presence of the parasite - a single infected individual is enough - and absence - that requires the whole species to be absent (Chase et al., 2019). Reliance on this binary matrix defined at the regional level also prevented the use of other statistics like the effective number of species (Chase and Knight, 2013) or individual based rarefaction (Chase et al., 2018) to better account for differences in scales (geographical and temporal) across the considered species sets. As a result, the significant species turnover across the recovery methods that we observed as other authors (Bellaw and Nielsen, 2020) may simply reflect scale differences between studies. We took care to correct for differences in sampling effort and restricted the analysis to the same geoclimatic conditions. But the wider diversity variance found for strongylid communities examined after horse deworming is also likely to reflect the higher abundance of this type of data relative to the necropsy-based communities as the more communities are sampled, the more likely that species recovery will vary (Chase et al., 2019).

Overall, our results highlighted a small group of abundant and widespread species *C. nassatus, C. longibursatus* and *C. catinatum* that clearly separated from other species. Of note, these species were consistently associated with drug resistance reports suggesting their phenology may have favoured higher fitness towards the use of modern anthelmintics (van Doorn et al., 2014). It is unclear however what phenotypic trait underpins their higher abundance and prevalence in horse populations. Their respective fecundities estimated from the number of eggs found in female worms *utero* was among the lowest suggesting their dispersal is unlikely to define a better ability to colonize their hosts (Sallé et al., 2018).

In addition, analysis of horse-based data expanded our understanding of strongylid communities. First, we found limited correlation between parasite egg excretion, a trait used to monitor infection in horses, and the strongylid community diversity. This was against our hypothesis that more permissive hosts would tolerate higher parasite burden, thereby increasing the community species richness. This lack of linear relationship may mirror variations in respective strongylid species fecundities (Kuzmina et al., 2012) and reflect possible density dependence effects that are known to occur in trichostrongylid species infecting ruminants (Bishop and Stear, 2001).

Second, we found that overall diversity was mostly partitioned across within-host richness and species turnover across horse age groups, and to a lesser-extent between farm variation. Species richness is conditional upon availability of host resources, leading to higher richness in larger animal species or for higher host density (Kamiya et al., 2014). Richness also depends on environment saturation in infective larvae (Shmida and Wilson, 1985). The latter two aspects underscore the importance of pasture hygiene and limited stocking density for the management of strongylid infection in grazing livestock. Of note, environmental saturation is usually expected to erase species turnover among hosts (Vannette and Fukami, 2017) but we identified a significant turnover between age groups. This turnover remains to be fully dissected to disentangle between the respective effects of selection applied through the mounting of an effective immune response as horses age (Debeffe et al., 2016; Kornaś et al., 2015; Relf et al., 2013; Wood et al., 2013) or the drift arising from putative competitive processes between species. To this respect, the increased species turnover across age groups may arise from priority effects between strongylid species that would counterbalance the effect of environmental saturation in eggs and infective larvae (Vannette and Fukami, 2017). Such priority effect would in turn yield a network of positive co-occurrence between species as found for mixed infection of rodents by malaria species (Ramiro et al., 2016).

Using data with anatomical niche resolution, we performed a first investigation of co-occurrence between strongylid species. This analysis did not reveal any negative effects thereby corroborating past results obtained after horse deworming in Poland conditions (Sallé et al., 2018). Of note, a few co-occurrences were dependent on associated covariates including the anatomical niche or horse age that may define optimal environmental conditions shared across parasite species. However, a core network of positive interactions remained after accounting for environmental variations between species from four distinct genera, namely *Cyathostomum (C. catinatum), Cylicocyclus* (*C. nassatus* and *C. insigne*), *Cylicostephanus (C. minutus)* and *Coronocyclus* (*C. coronatus* and *C. labratus*). This would be compatible with facilitative relationships between more divergent species that should be less prone to resource competition (MacArthur and Pianka, 1966) although co-existence of the *Cylicocyclus* and *Coronocyclus* members depart from this expectation. While the complexity of underpinning biological processes may obscure the observed co-occurrences patterns (Blanchet et al., 2020), these interactions were found across different parasite communities and with the lowest possible resolution thereby supporting their biological meanings (Behnke, 2008). However, covariates like past infections and treatment history were not accounted for and may contribute to this network. Besides, the contribution of the resident gut microbiota may play a role in the definition of this particular structuring as significant interactions between microbial and helminth communities are known to occur in horses (Clark et al., 2018; Peachey et al., 2019; Walshe et al., 2020).

Altogether, priority effects between equine strongylid species in young horses may contribute to enhance the species turnover found between age groups and to define the network of positive co-occurrences between the most dominant genera. The simultaneous monitoring of both the microbial and parasite communities using the latest barcoding approaches (Poissant et al., 2020) should provide valuable contributions to challenge these hypotheses.

## Acknowledgements

MBo and this work was supported by the Institut Français du Cheval et de l’Équitation (IFCE) and Fonds Éperon grant.

