## Supplementary Information for "Species turnover between age groups of horses and positive network of co-occurrences define the structure of horse strongylid communities: a meta-analysis"

**Supplementary table 1. List of studies considered in the meta-analysis with associated meta-data. 2**

**Supplementary table 2: Table of average abundances and prevalences for each strongyles species. 2**

**Supplementary Figure 1. Relationship between Faecal Egg Count and strongyle species diversity 3**

**Supplementary Figure 2. Principal Coordinate Analysis of community diversity gathered under continental European conditions. 4**

**Supplementary Figure 3. Distribution of species presence/absence rate across recovery methods 5**

**Supplementary Figure 4. Distribution of species presence/absence rate across geoclimatic areas 6**

**Supplementary Figure 5. Variance partitioning of species occurrence from positive abundance data across considered covariates 7**

**Supplementary Figure 6. Variance partitioning of species occurrence across considered covariates following presence-absence modeling 8**

**Supplementary table 1. List of studies considered in the meta-analysis with associated meta-data.**

See corresponding attached csv file.

**Supplementary table 2: Table of average abundances and prevalences for each strongyles species.**

“Species_code” corresponds to the three-letter code associated with each species. “Species” correspond to the full name (genus and species). “Prevalence1” and “abundance” correspond to the relative abundance and mean prevalence calculated with the data collected in the studies. Finally, “Prevalence2” is the prevalence we estimated with our presence and absence matrix.

**Supplementary Figure 1. Relationship between Faecal Egg Count and strongyle species diversity**

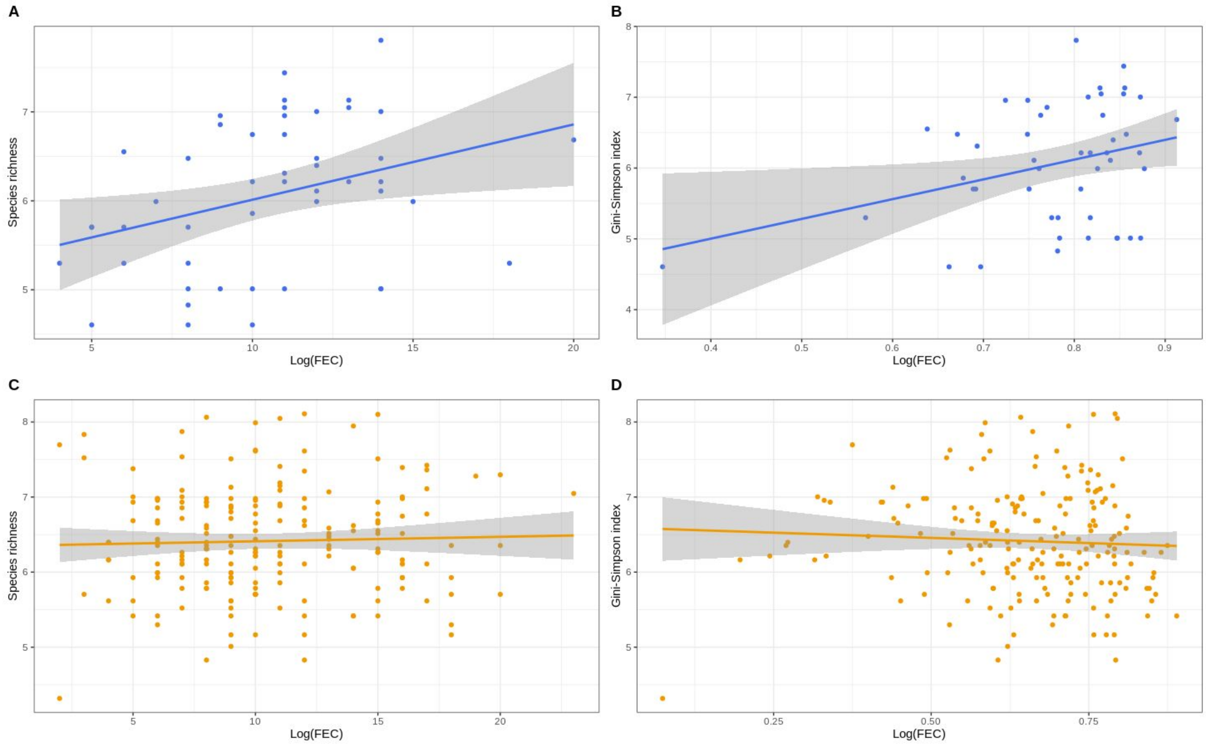


Species richness (left panels) or the Gini-Simpson index is plotted against log-transformed Faecal Egg Count measured in 48 horses from Poland (upper panels) and 197 horses from Ukraine (lower panels).

**Supplementary Figure 2. Principal Coordinate Analysis of community diversity gathered under continental European conditions.**

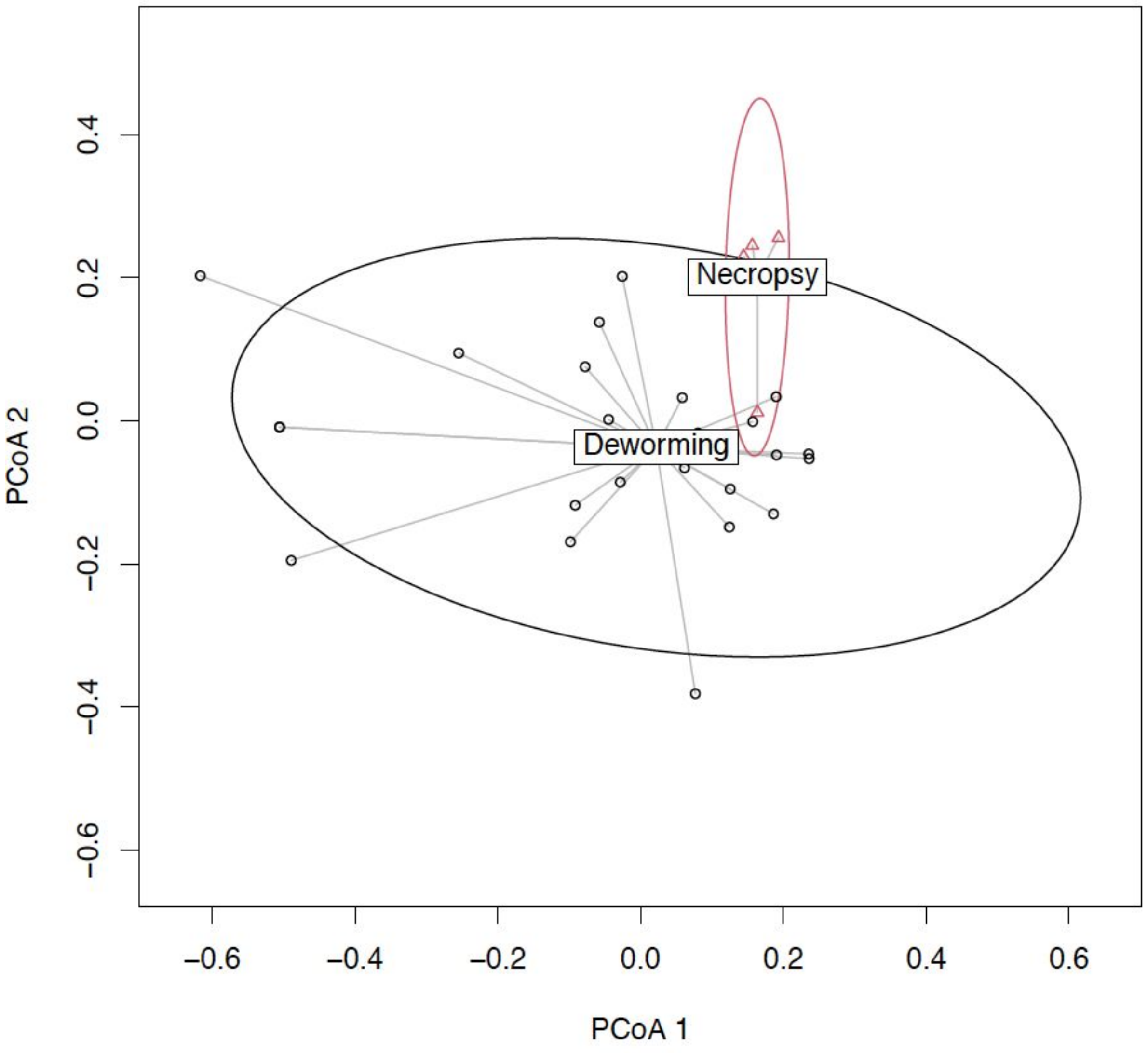


This figure illustrates the multivariate homogeneity of between group dispersions for studies using either worm collection upon horse necropsy (red) or collection after deworming (in black). Ellipses indicate associated 95% confidence intervals.

**Supplementary Figure 3. Distribution of species presence/absence rate across recovery methods**

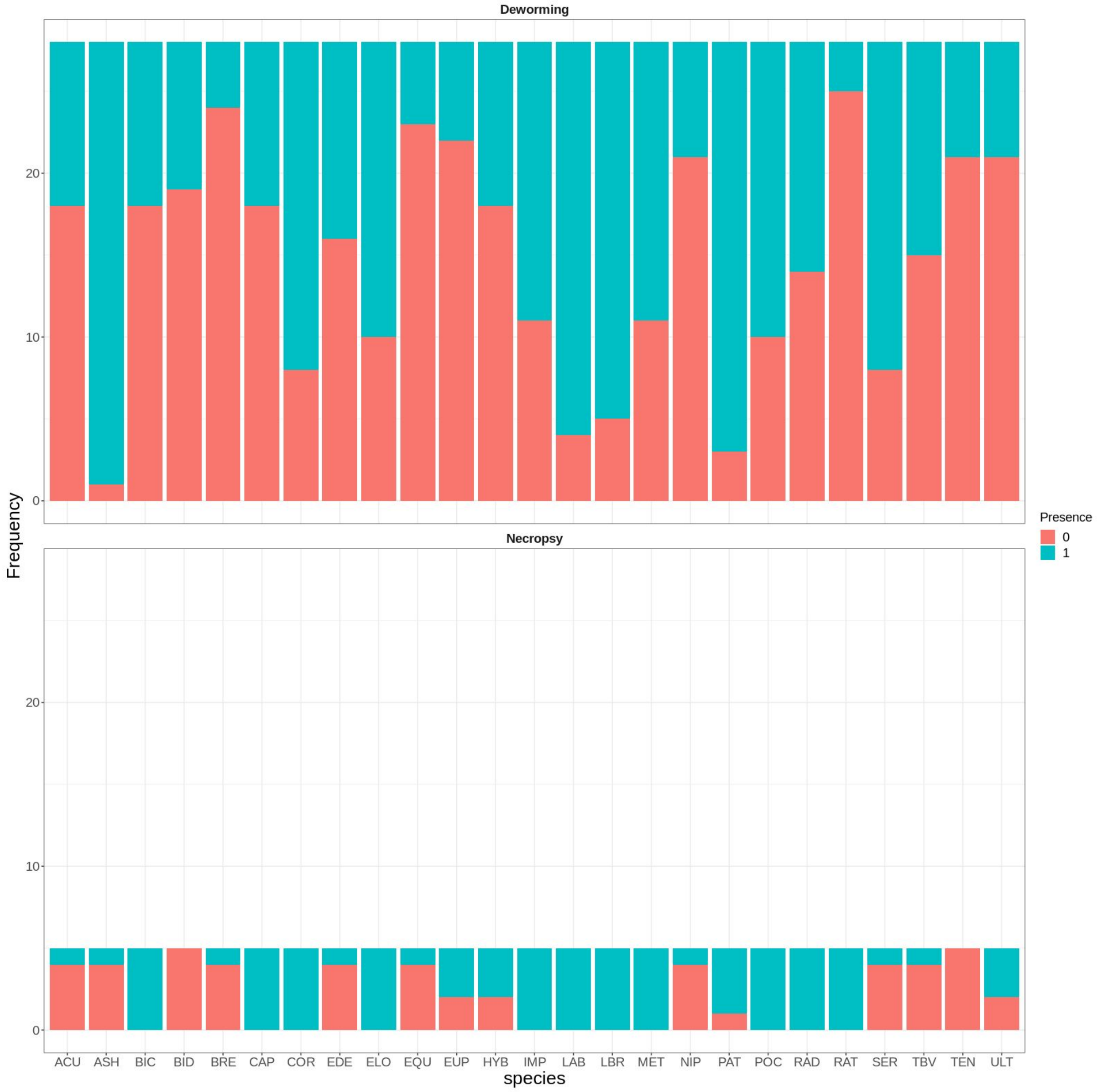


The bar chart illustrates each species presence (green) or absence (red) across published studies performed under European continental area either following deworming (upper panel) or necropsy (lower panel). Species with too extreme prevalence (i.e. less than 10% or more than 90% were not considered for this analysis).

**Supplementary Figure 4. Distribution of species presence/absence rate across geoclimatic areas**

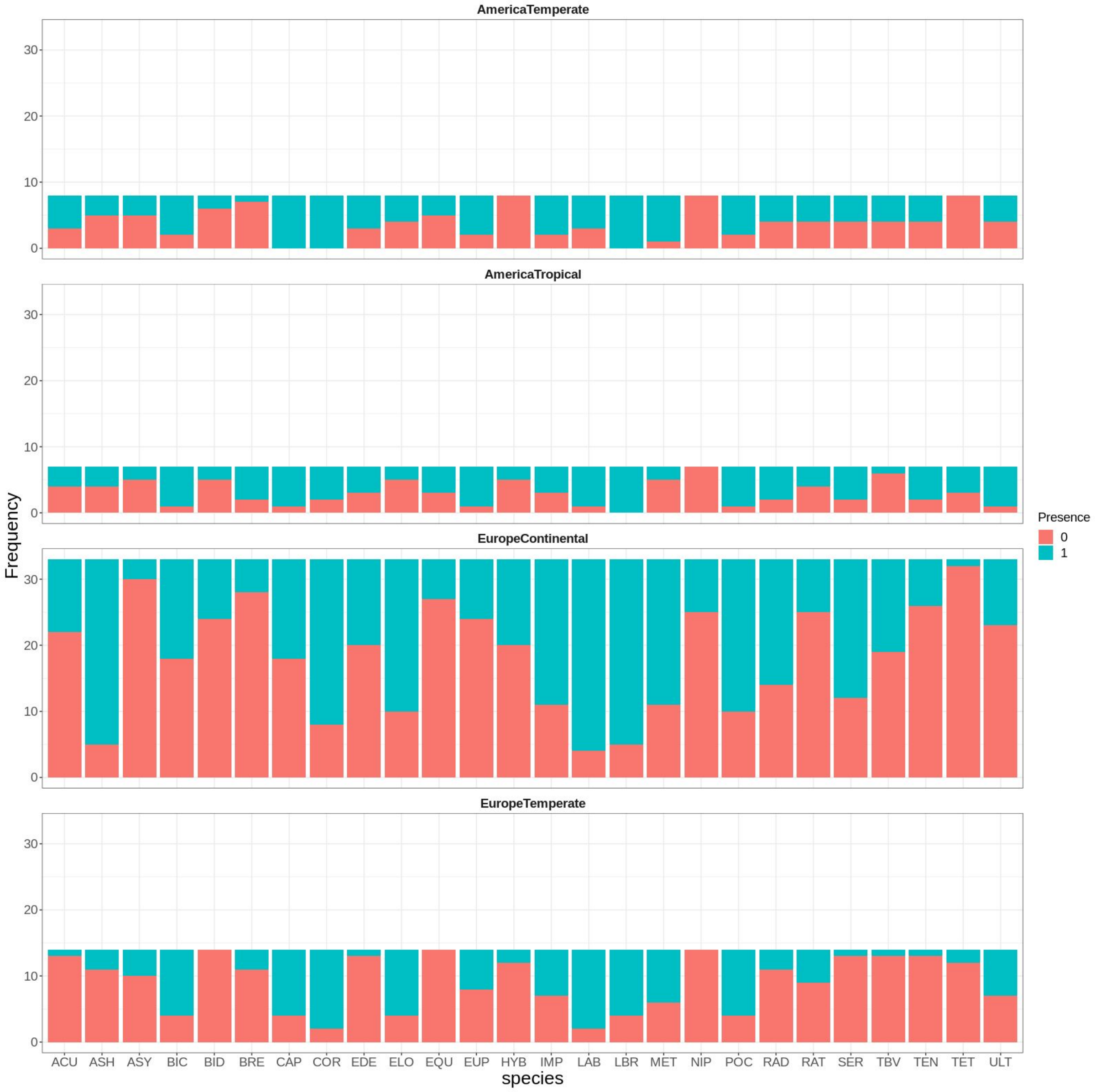


The bar chart illustrates each species presence (green) or absence (red) across geo-climatic conditions in studies collecting worms after necropsy. Species with too extreme prevalence (i.e. less than 10% or more than 90% were not considered for this analysis).

**Supplementary Figure 5. Variance partitioning of species occurrence from positive abundance data across considered covariates**

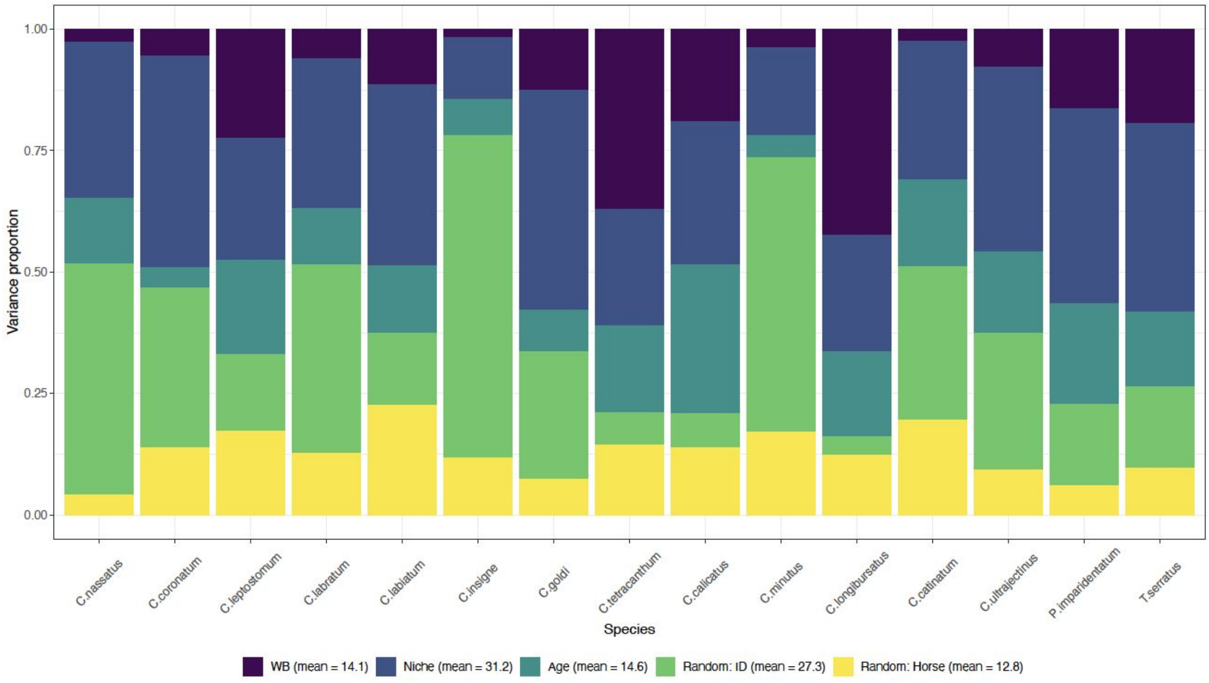


For each species, the relative proportion of variance explained by each fixed or random covariate is illustrated. This is based on the analysis of species positive abundances.

**Supplementary Figure 6. Variance partitioning of species occurrence across considered covariates following presence-absence modeling**

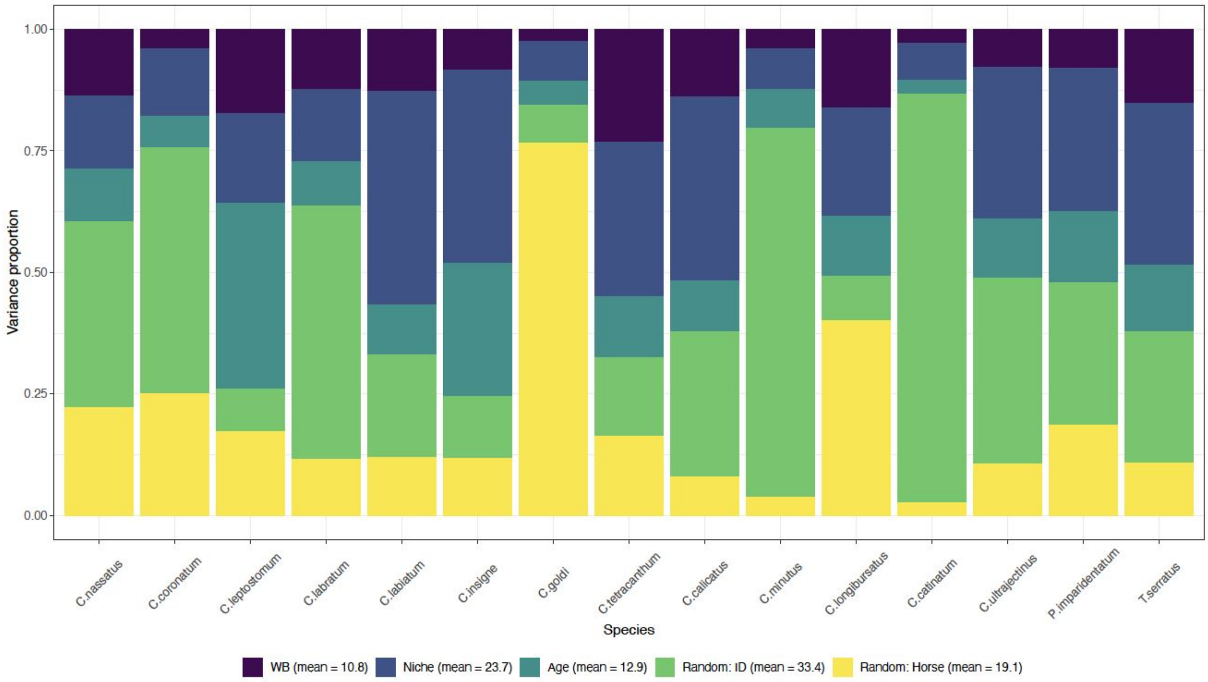


For each species, the relative proportion of variance explained by each fixed or random covariate is illustrated.
